# Presenilin-1 controls glucose metabolism and identity of pancreatic beta cells

**DOI:** 10.1101/2025.09.05.674426

**Authors:** Zhanat Koshenov, Sandra Postic, Gabriela Schosiwohl, Savina van Amsterdam, Victoria Hois-Zelinka, Furkan E. Oflaz, Rene Rost, Oleksandra Tiapko, Benjamin Gottschalk, Martin Hirtl, Olaf G. Bachkoenig, Aigerim Koshenova, Yusuf C. Erdogan, Adlet Sagintayev, Anita Krnjic, Johannes U. Pfabe, Srdjan Sarikas, Bernhard Hochreiter, Juergen Gindlhuber, Matthias Schittmayer, Jelena Tadic, Barbara Ehall, Frank Madeo, Tobias Madl, Roland Malli, Ruth Birner-Gruenberger, Thomas Pieber, Tobias Eisenberg, Marjan Slak Rupnik, Wolfgang F. Graier

## Abstract

Presenilin 1 is an endoplasmic reticulum protein, most known for its role in pathogenesis of familial Alzheimer’s Disease (AD). Presenilin 1 has been attributed roles in intracellular calcium homeostasis in the brain, as well as in the pancreatic beta cells, where it has been shown to be fundamental for glucose-induced insulin secretion. Functional similarity of presenilin 1 in regulation of intracellular calcium homeostasis in the brain and pancreas prompted us to investigate a prevalent assumption that associates AD and diabetes mellitus. By examining pancreatic islets from AD model mice, we have found deficits in initial phase of glucose-induced calcium signaling and insulin secretion. Furthermore, these transgenic mice showed a tendency towards reduced expression of mature beta cell markers, which was even more pronounced in islets and beta cell lines with a transient knock down of presenilin 1. We demonstrate here that presenilin 1 controls beta cell glycolysis by regulating sub-cellular calcium homeostasis and, in doing so, contributes to preservation of beta cell identity.

## Introduction

Pancreatic beta cells function as metabolic sensors of the body. They detect and respond to elevated blood glucose by a series of metabolic events culminating in insulin secretion, which dictates whole body glucose metabolism. The molecular machinery behind this robust response is tightly controlled, with Ca^2+^ ions playing the key regulatory roles^1^. Following glucose uptake by plasma membrane glucose transporters (mainly GLUT 2 in rodents and GLUT 1 and 3 in humans)^2^, glucose is metabolized by beta cells to generate ATP. While most of the ATP is generated in the mitochondria, glycolysis plays a major role in beta cell metabolism^3^. Both mitochondrial oxidative phosphorylation (OXPHOS) and cytosolic glycolysis can be regulated by Ca^2+^ ions^1,4,5^, adding complexity to bioenergetic control of these highly specialized glucose sensor cells.

The molecular machinery of beta cells’ glucose metabolism is highly conserved and is part of beta cell identity^6^. At the genetic level, beta cell identity is maintained by activity of key transcription factors, such as pancreatic and duodenal homeobox 1 (PDX1), MAF BZIP Transcription Factor A (MafA), and NK6 homeobox 1 (NKX6.1)^6–8^. Loss of beta cell identity is associated with the development of type 1 and 2 diabetes^7–10^. Interestingly, metabolic control of cell fate and identity has recently emerged as a prominent topic across many fields of biology. Thus, identification of novel genetic and metabolic control mechanisms of metabolism and identity of beta cells can have a fundamental impact on our understanding of beta cell function and can provide new therapeutic targets against diabetes.

There is a large body of literature associating Alzheimer’s disease (AD) with diabetes and often calling AD a diabetes of the brain^11–14^. Indeed, there are particular similarities between the diseases, especially when it comes to glucose metabolism and Ca^2+^ signaling^11,15^. Our previous work has shown that presenilin 1 (PS1) plays a fundamental role in establishing an endoplasmic reticulum (ER) Ca^2+^ leak directed towards mitochondria that is needed for proper glucose responsiveness of beta cells^16,17^. Such role of PS1 in pancreatic beta cells displays parallels in neurons, where it is also responsible for ER Ca^2+^ leak, and loss or mutation of PS1 leads to neurodegenerative disorders, including familial AD^15^. We undertook the current study to investigate whether PS1 could provide a mechanistic link between the two devastating disorders of the brain and pancreas. Since more is known about the role of PS1 in the brain, we focused on the impact of loss or mutation of PS1 in the pancreas to identify its detailed mechanism of action in this organ. As a result, we uncovered mechanistic details of PS1 mediated control of beta cell metabolism and revealed a novel role of PS1 as a regulator of beta cell identity.

## Results

### Genetic driver of the Familial Alzheimer’s Disease causes impaired glucose responsiveness and identity loss in pancreatic islets

We have previously shown that PS1 establishes an ER Ca^2+^ leak in pancreatic beta cells that is important for glucose-induced insulin secretion^17^. Since PS1 mutations are a major cause of familial AD^15,18^ and given a large amount of published work claiming high correlation between Alzheimer’s disease and diabetes^11–14,19^, we wondered if drivers of AD could cause dysfunction in pancreatic beta cells. Hence, we have used a triple transgenic (3xTg) AD mouse model that have a brain-restricted expression of human amyloid precursor protein with the Swedish mutation (APPSwe) and tubulin-associated unit (Tau) with P301L mutation, while the human PS1 gene with M146V mutation is expressed in the whole body, including the pancreas^20,21^. Glucose-stimulated Ca^2+^ oscillations were delayed and reduced in pancreatic organ slices from 3xTg mice (Figure 1 a-b), similar to our previous^17^ and current findings with transient PS1 KD (Figure S1 a-e). In line with delayed onset of glucose stimulated Ca^2+^ oscillations, islets from the AD model mice showed decreased insulin secretion during the first 5 minutes of glucose stimulation (Figure 1 d). There was no difference in total islet insulin content or insulin secretion measured over a prolonged time (30 minutes) (Figure 1 e, f), similar to our previous findings with PS1 KD in pancreatic islets^17^. These results point to a specific role of PS1 in the initial steps of glucose response in pancreatic beta cells and might offer an explanation for why there were only small changes in blood glucose level of *ad libitum* fed AD model mice (Figure S1 f).

**Figure 1.**
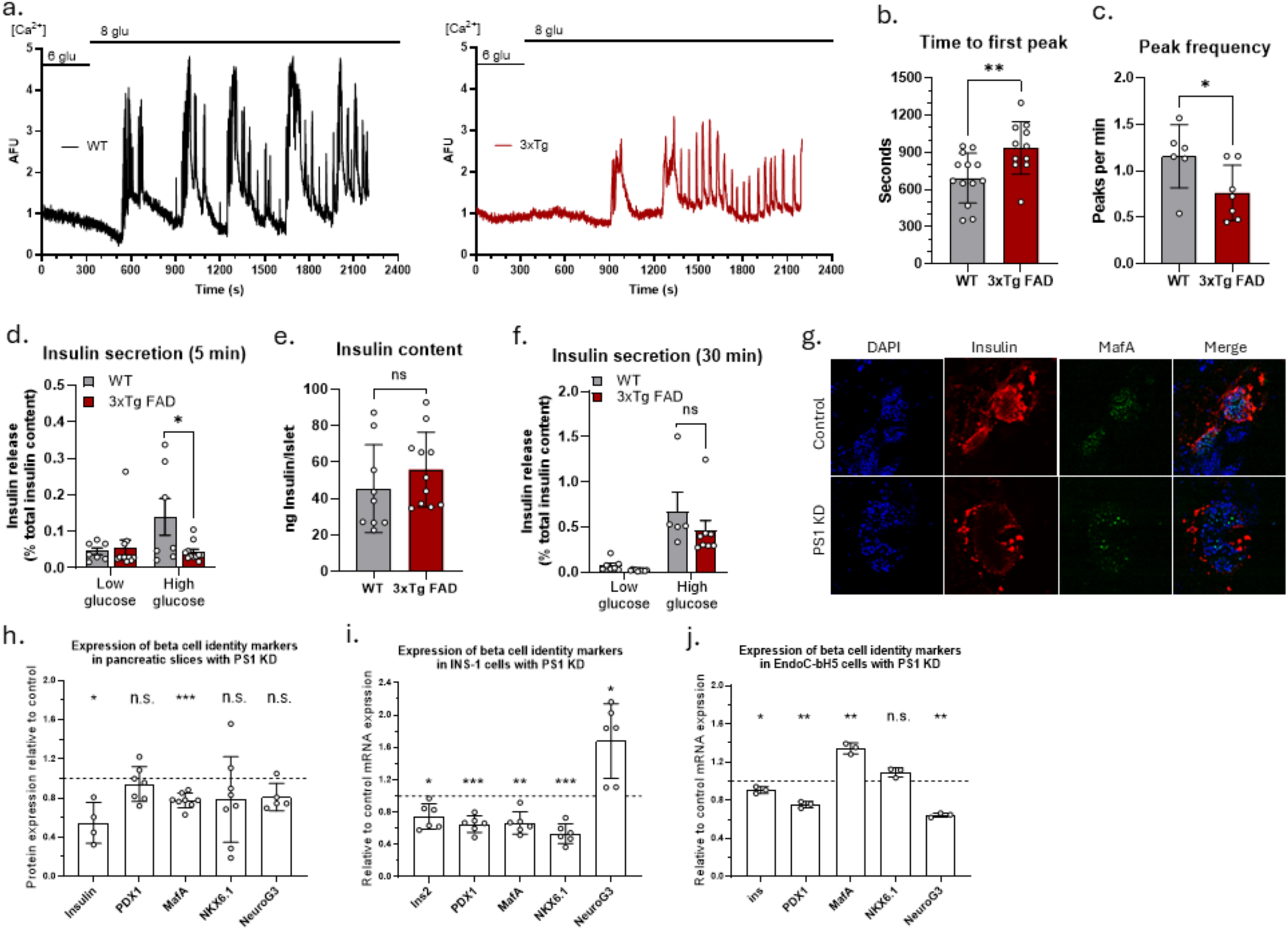
PS1 is important for glucose sensitivity and identity of pancreatic beta cells. **a,** representative traces showing cytosolic Ca^2+^ oscillations in response to elevation of extracellular glucose in beta cells from pancreatic slices of WT (left) and 3xTg AD model (right) animals. **b,** statistical analysis of experiments as shown in **a,** bar graphs (+/-SD) show a time delay between the addition of 8 mM glucose and the onset of Ca^2+^ oscillations (n=11-13). **c,** bar graphs (+/-SD) show the cytosolic Ca^2+^ peak frequency for experiments as shown in **a** (n=6-7). **d,** insulin secretion (+/-SD) during the first 5 minutes of glucose stimulation by pancreatic islets from WT and 3xTg AD model mice (n=7-11). **e**, total insulin content (+/-SD) in islets from WT and 3xTg FAD mice (n=7-11). **f**, extended insulin secretion (+/-SD) during the first 30 minutes of glucose stimulation by pancreatic islets from WT and 3xTg AD model mice (n=7-11). **g**, representative images of control and PS1KD pancreatic organ slices from WT mice, stained for beta cell identity markers, insulin and MafA; **h**, analysis of expression of beta cell identity markers in control and PS1 KD pancreatic slices as shown in **g**; values are normalized to respective average control expression (n=4-8). Expression of beta cell identity markers in control and PS1 KD INS1 (**i**, n=6) and EndoC-bH5 (**j**, n=3) cells; values are normalized to respective average WT expression. Unpaired t-test (**b-f**), paired t-test (**g-k**), n.s. p > 0.05, * p < 0.05, ** p < 0.01, *** p < 0.001)

Glucose metabolism plays a central role in the identity of pancreatic beta cells^6^. As our experiments indicate an involvement of PS1 in glucose responsiveness of beta cells, we tested if it also influences beta cell identity. For that, we analyzed islets from 3xTg mice for markers of mature beta cells and found that AD model mice had slightly reduced expression of key identity markers, such as *Pdx1*, *MafA,* and *Nkx6.1*, while the expression of insulin and *NeuroG3*, a pancreatic islet progenitor marker, were unaffected (Figure S1 a). Identity changes were more pronounced in mouse islets and cultured human and rat beta cells with transient KD of PS1 (Figure 1 g-j, Figure S1 g-i), suggesting an important role for PS1 in preservation of beta cell identity. Interestingly, each beta cell model had a slightly different signature of identity loss in response to transient PS1 KD, with the most affected model being INS-1 cell line. For that reason and given the ease of manipulation, we used this cell line to further investigate the mechanism of PS1’s involvement in preservation of beta cell identity.

### Presenilin 1 regulates the activity of transcription factors sensitive to oscillatory Ca^2+^ signals

Given PS1’s role in ER Ca^2+^ leak^15–17^ and Ca^2+^ signaling during glucose stimulation, we investigated the activities of various Ca^2+^ sensitive regulatory proteins and transcription factors that could have been controlled by PS1 for preservation of beta cell identity. We found that PS1 KD in INS-1 cells did not affect cAMP response element binding protein (CREB) and calcium-calmodulin (CaM)-dependent protein kinase II (CaMKII) activities (Figure S2 a-d). In contrast, PS1 depletion increased the nuclear translocation of several NFAT family members (NFATc2 and NFATc3), while showing no effect on NFATc1 (Figure 2 a, b). Since we have observed reduced glucose-induced Ca^2+^ signaling in PS1 KD beta cells, the increased NFAT nuclear translocation, which is triggered by Ca^2+^ signals, came as a surprise.

**Figure 2.**
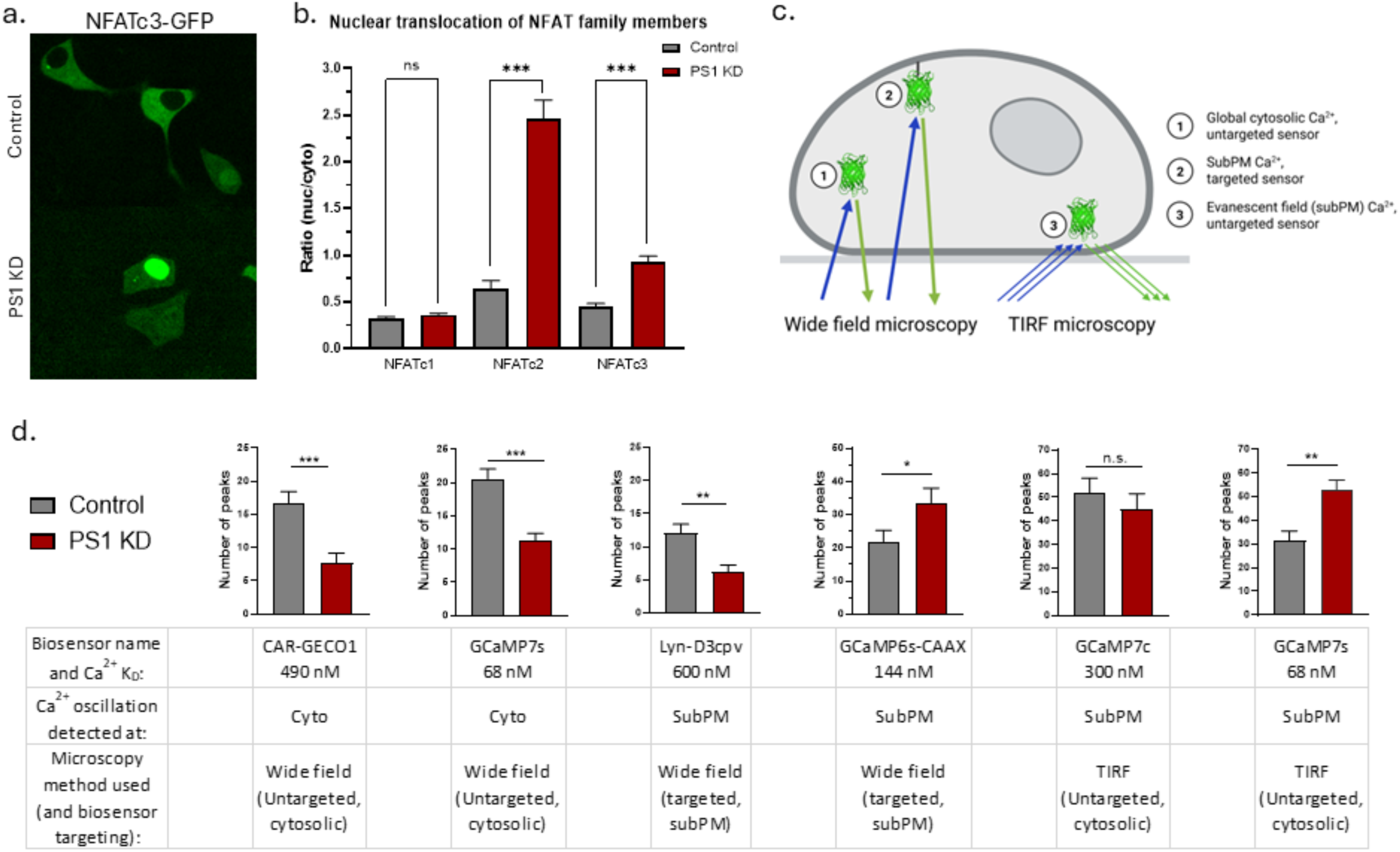
Presenilin-1 contributes to resting sub-plasma membrane Ca^2+^ signaling. **a,** representative images of INS-1 cells overexpressing NFATc3-GFP construct to monitor its nuclear localization. **b,** analysis of NFATc1-3 nuclear/cytosolic localization ratio (control: NFATc1 n=128, NFATc2 n=66, NFATc3 n=128; PS1 KD: NFATc1 n=112, NFATc2 n=76, NFATc3 n=141). **c,** graphical representation of the experimental set up used for investigation of sub-cellular Ca2+ oscillation pattern in INS-1 cells. **d**, basal global cytosolic and subPM Ca^2+^ oscillations measured with different techniques using various biosensors (shown in **c** and outlined in a table below bar the graphs); (Wide field: control n=41-72, PS1 KD n=47-59; TIRF: control n=5-14, PS1 KD n=4-13). Unpaired t-test, n.s. p > 0.05, * p < 0.05, ** p < 0.01, *** p < 0.001)

To explore the reason for such diverse conduct of the transcriptional activities of NFAT during KD of PS1, we investigated the impact of PS1 depletion on sub-cellular Ca^2+^ oscillation patterns in unstimulated INS-1 cells. Therefore, basal Ca^2+^ oscillations in the global cytosol or the sub-plasma membrane (subPM) region were recorded using Ca^2+^ sensors with low Ca^2+^ affinity (K_D_∼490 nM for CAR-GECO1^22^, and ∼600 nM for Lyn-D3cpv^23^), medium Ca^2+^ affinity (K_D_∼300 nM for GCaMP7c^24^), and high Ca^2+^ affinity (K_D_∼144 nM for GCaMP6s-CAAX^25^ and ∼68 nM for GCaMP7s^24^). For the subPM Ca^2+^ imaging, the sensors were specifically targeted to subPM (Lyn-D3cpv and CAAX-GCaMP6f) and imaged with wide-field microscopy, or the sensors were untargeted (GCaMP7s and GCaMP6f) but imaged in close proximity to the basal plasma membrane with total internal reflection microscopy (TIRF) (Figure 2 c). These experiments revealed a differential contribution of PS1 to sub-cellular Ca^2+^ oscillations (Figure 2 d). PS1 KD reduced global cytosolic Ca^2+^ oscillations, measurable with low- and high-affinity Ca^2+^ sensors. On the other hand, PS1 KD resulted in increased Ca^2+^ oscillations in the subPM region that were only detected with the high sensitivity Ca^2+^ sensors (CAAX-GCaMP6f, GCaMP7s), but not low affinity Ca^2+^ sensors (Lyn-D3cpv, GCaMP7c), suggesting an exclusive increase in low-amplitude Ca^2+^ oscillations. These divergent changes in subPM Ca^2+^ oscillations compared to those found in the cytosol could be responsible for the differential activities of NFAT family members.

To further corroborate these findings, we performed patch-clamp experiments on control and PS1 KD cells under the current clamp condition. We observed depolarization of the PM potential by PS1 KD, as well as an increased frequency of small-amplitude action potentials (AP) (Figure S2 e, f), which might reflect the increase in low amplitude Ca^2+^ oscillations in the subPM region.

### Loss of presenilin-1 leads to altered calcineurin activity, causing AMPK/mTORC1 imbalance

Our findings on altered sub-cellular Ca^2+^ oscillations and altered activation pattern of NFAT signaling point at a possible link to heterogeneous activation of calcineurin. It is known that calcineurin can be differentially regulated by sub-cellular Ca^2+^ signaling^26^, thus leading us to investigate its sub-cellular activation status. For that, we used calcineurin activation status sensor CaNARi^26^ in the cytosol (cyto CaNARi), targeted to the outer mitochondrial membrane facing the cytosol (OMM CaNARi), and the subPM region (subPM CaNARi) in INS-1 cells. We observed differential impacts of PS1 KD on the activation status of calcineurin. PS1 KD had no effect on calcineurin activity in the cytosol but resulted in increased activity in the sub-PM region and reduced activity at the OMM (Figure 3 a). Increased calcineurin activation at the sub-PM region likely stems from increased Ca^2+^ oscillations at the same microenvironment, since addition of verapamil, an inhibitor of voltage gated Ca^2+^ channels normalized the calcineurin activities of control and PS1 KD cells at sub-PM region.

**Figure 3.**
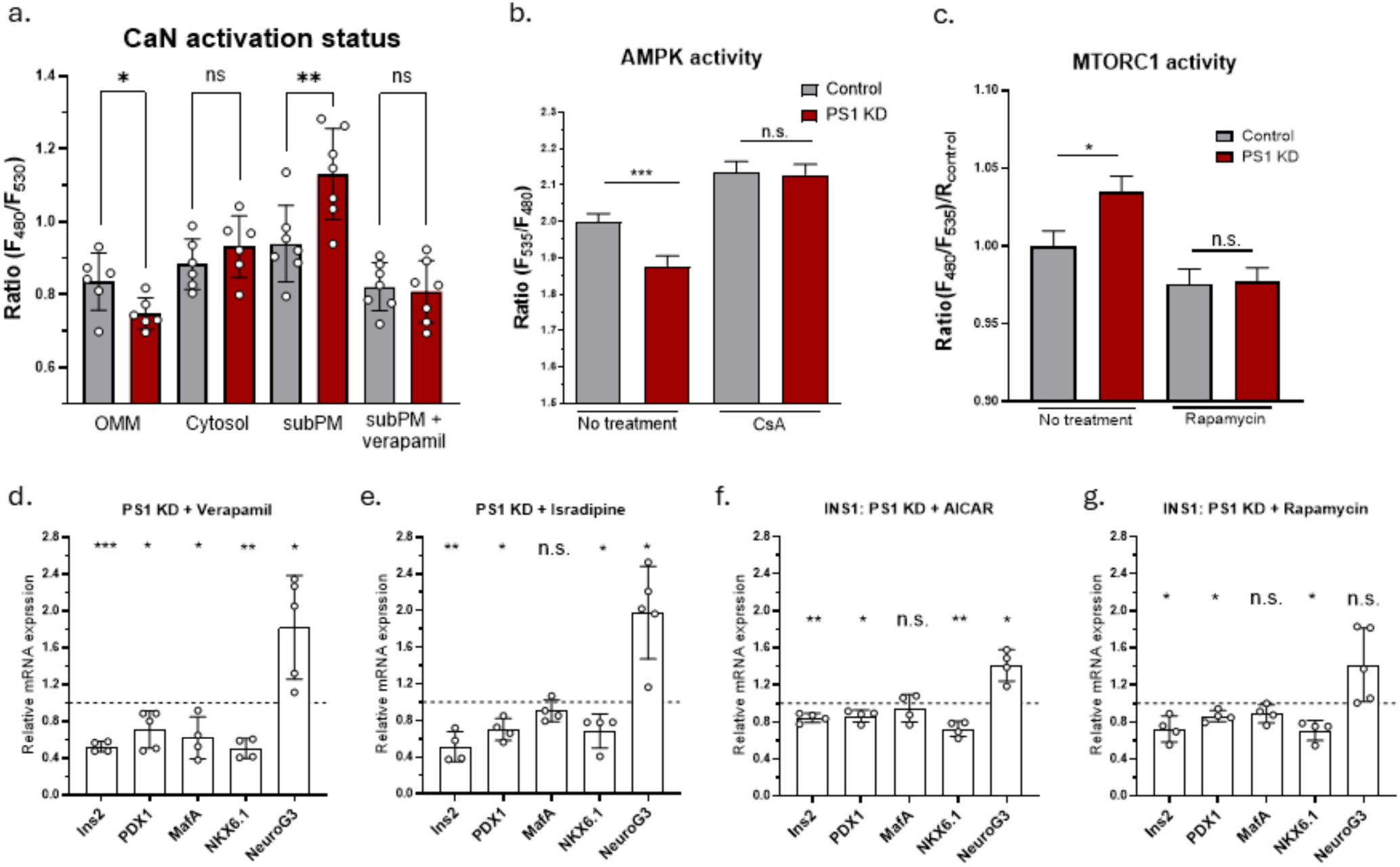
Presenilin-1 regulates spatial calcineurin activity and exerts control over AMPK-mTOR pathways. **a,** calcineurin activation status at different subcellular compartments measured with CaNARi and shown as mean emission ratio (+/-SD) for control and PS1 KD INS-1 cells (n=6-7). **b,** AMPK activity measured with AMPKAR in INS-1 cells and shown as mean emission ratio (+/-SEM, control no treatment n=110, control CsA n=40, PS1 KD no treatment n=112, PS1 KD CsA n=41). **c,** mTORC1 activity measured with TORCAR in INS-1 cells and shown as mean emission ratio (+/-SEM, control no treatment n=51, control rapamycin n=40, PS1 KD no treatment n=51, PS1 KD rapamycin n=42). **d-g,** bar graphs show mean (+/-SD) mRNA expression of insulin, PDX1, MafA, NKX6.1 and Neurogenin-3 in PS1 KD INS-1 cells under various experimental conditions (**d**, verapamil, n=4-5; **e**, isradipine, n=4-5; **f**, AICAR, n=4; **g**, Rapamycin, n=4-5) normalized to respective mRNA expression in control cells (dotted line at Y=1) under matching experimental condition. Unpaired t-test (**a-c**), paired t-test (**d-g**), n.s. p > 0.05, * p < 0.05, ** p < 0.01, *** p < 0.001)

Besides its role in the activation of NFAT, calcineurin dephosphorylates and negatively regulates the 5’ AMP-activated protein kinase (AMPK)^27,28^, which is a well-established energy sensor in many cell types and plays a major role in beta cell development^29–32^. In line with increased calcineurin activation at subPM region, PS1 KD resulted in reduced AMPK activity in INS-1 cells. This increase was indeed calcineurin dependent, as Cyclosporin A (CsA), a known calcineurin inhibitor, increased AMPK activity in both control and PS1 KD cells to comparable levels (Figure 3 b). Furthermore, the mammalian target of the rapamycin complex 1 (mTORC1) is known to be involved in beta cell identity downstream of AMPK, where increased mTORC1 activity is linked to loss of beta cell identity^33–35^. Therefore, we next tested the impact of PS1 KD on the mTORC1 pathway and found that PS1 KD results in increased mTORC1 activity (Figure 3 c).

This PS1 and calcineurin dependent reversal in AMPK and mTORC1 activities could be responsible for the loss of beta cell identity. To test this claim, we have attempted to rescue loss of beta cell identity in PS1 KD cells by activating AMPK or inhibiting mTORC1, with AICAR and rapamycin, respectively. Additionally, we have used verapamil and isradipine, two voltage gated Ca^2+^ channel blockers, to normalize calcineurin activity. To our surprise, none of these approaches fully rescued the expression of mature beta cell markers (Figure 3 d-g), suggesting that PS1 likely has additional mechanisms to influence beta cell identity.

### Presenilin-1 is needed for glycolytic flux in pancreatic beta cells

Taking into account reduced calcineurin activation on the mitochondrial surface, reduced ER Ca^2+^ leak towards mitochondria^16^ and reduced cytosolic Ca^2+^ oscillations in PS1 KD cells, we wondered if loss of PS1 results in defects in mitochondrial metabolism that could explain reduced glucose sensitivity and loss of identity phenotypes. To test this hypothesis, we measured Ca^2+^ signals in response to stimulation with pyruvate, instead of glucose, thus, directly feeding mitochondrial energy metabolism. This experiment showed no difference in onset and number of Ca^2+^ peaks between control and PS1 KD INS-1 cells (Figure 4 a, b), suggesting that mitochondria are not responsible for reduced glucose sensitivity in beta cells lacking PS1. Additionally, this experiment suggests that PS1 might have a role in glycolysis.

**Figure 4.**
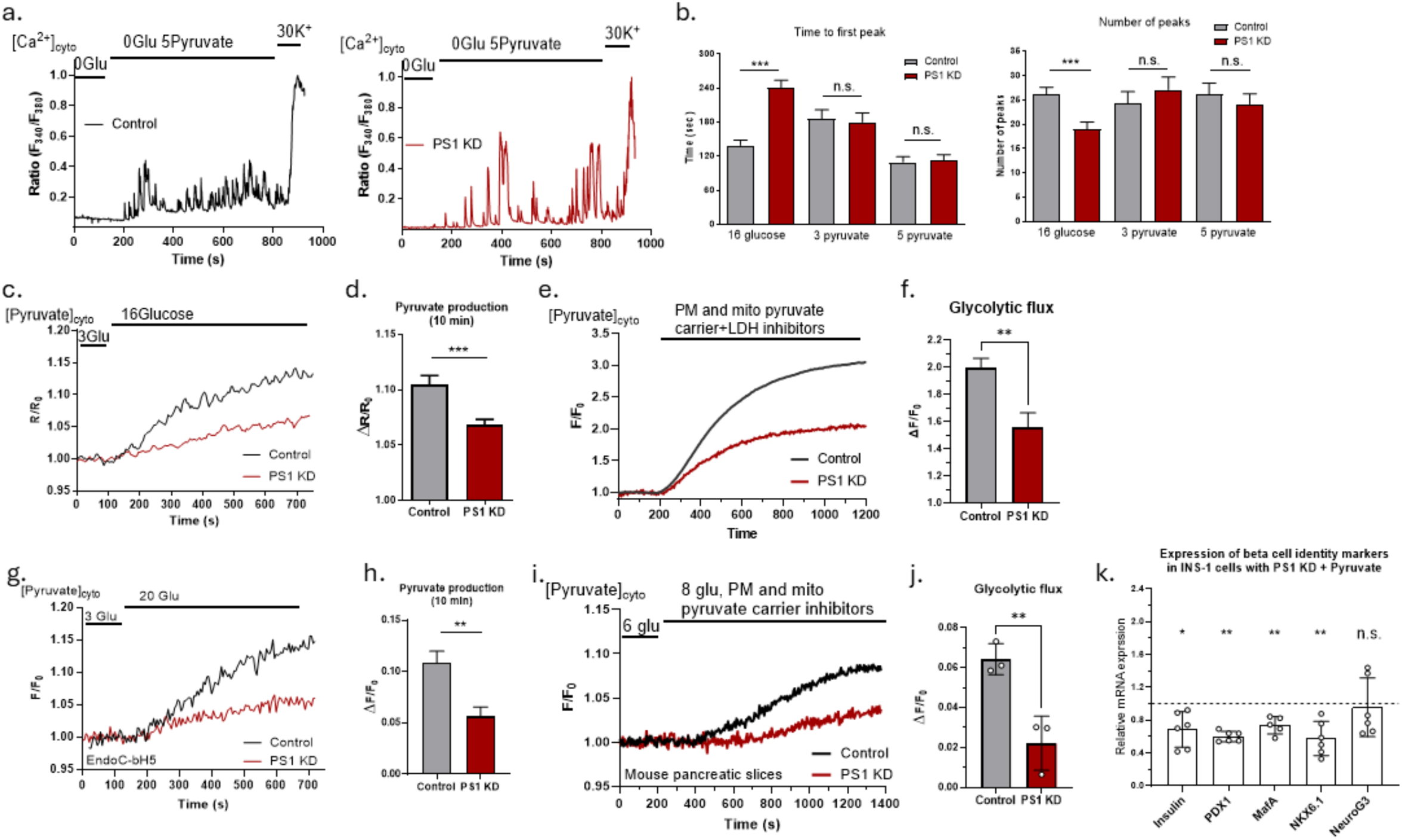
Presenilin-1 regulates beta cell metabolism by controlling glycolytic flux. **a,** representative FURA-2 AM traces depicting cytosolic Ca^2+^ oscillations in response to 5 mM pyruvate in control (black traces) and PS1 KD (red traces) INS-1 cells. **b**, statistical analysis of experiments as shown in **b,** bar graphs (+/-SEM) on the left panel show time delay between the addition of 3-5 mM pyruvate and the onset of Ca^2+^ oscillations; bar graphs (+/-SEM) on the right panel show the number of Ca^2+^ oscillations/peaks (control, 3 mM pyruvate, n=78; PS1 KD, 3 mM pyruvate, n=93; control, 5 mM pyruvate, n=81; PS1 KD, 5 mM pyruvate, n=82). **c,** representative Pyronic traces depicting cytosolic pyruvate dynamics in response to increased extracellular glucose in control (black) and PS1 KD (red) INS-1 cells. **d,** statistical analysis of the experiment shown in **c**, bar graphs (+/-SEM) show change in cytosolic pyruvate after 10 min in 16 mM glucose (control, n=85, PS1 KD, n=112). **e,** representative PyronicSF traces depicting glycolytic flux experiment in control (black) and PS1 KD (red) INS-1 cells. **f,** quantification of the basal glycolytic flux (+/-SEM) as a change in cytosolic pyruvate after 15 min perfusion with the inhibitor mix (control, n=33, PS1 KD, n=29). **g,** representative PyronicSF traces depicting cytosolic pyruvate dynamics in response to increased extracellular glucose in control (black) and PS1 KD (red) EndoC-bH5 cells. **h,** statistical analysis of the experiment shown in **g**, bar graphs (+/-SEM) show change in cytosolic pyruvate after 10 min in 16 mM glucose (control, n=11, PS1 KD, n=12). **i,** representative PyronicSF traces in response to increased extracellular glucose and application of pyruvate carrier inhibitors in control (black) and PS1 KD (red) beta cells in primary pancreatic slices. **j,** quantification of the stimulated glycolytic flux (+/-SEM) as a change in cytosolic pyruvate after 15 min in 8 mM glucose and inhibitor mix (control, n=3, PS1 KD, n=3). **k**, expression of beta cell identity markers in PS1 KD INS-1 cells supplemented with pyruvate; values are normalized to respective average WT expression under the same experimental condition, n=6. Unpaired t-test (all, except **k**), paired t-test (**k**), n.s. p > 0.05, * p < 0.05, ** p < 0.01, *** p < 0.001)

In order to test glycolytic activity, we measured cytosolic pyruvate production after the elevation of extracellular glucose in INS-1 cells using pyruvate biosensors^36,37^. PS1 KD did not affect basal pyruvate level at low glucose (3 mM) (Figure S3 a), but it reduced pyruvate production upon glucose stimulation (16 mM) (Figure 4 c, d). To test the possibility of hampered glucose uptake, we measured cytosolic glucose accumulation in response to increasing extracellular glucose concentration. We found no impact of PS1 loss on glucose uptake but rather saw an increased accumulation of glucose (Figure S3 b, c).

In addition to pyruvate production upon glucose stimulation, we measured basal glycolytic flux at fixed glucose concentration by acute inhibition of pyruvate exit routes by inhibiting mitochondrial pyruvate carrier (MPC) using 10 µM UK-5099^38^, pyruvate and lactate extrusion by monocarboxylate transporter (MCT) using 1 µM AR-C155858^39^, and lactate dehydrogenase (LDH) using 1 µM GSK-2837808A^40^. The resulting accumulation of pyruvate upon inhibiting its exit routes was assessed as steady-state glycolytic flux. PS1 KD reduced glycolytic flux in INS-1 cells by about 50% (Figure 4 e, f). These experiments showed that PS1 indeed has a role in glycolytic flux, which could explain the reason for reduced glucose sensitivity of beta cells and islets lacking functional PS1. To further corroborate these findings, we measured glycolytic activity in human cultured beta cells (EndoC-bH5) and mouse islets with transient PS1 KD. Both these beta cell models supported our findings in INS-1 cells and demonstrated reduced glycolytic activity (Figure 4 g-j).

Having established the involvement of PS1 in beta cell glycolysis, we attempted to find out if this function is also important for the preservation of beta cell identity. To test this, we provided INS-1 cells with extra pyruvate to alleviate the impact of PS1 KD on reduced glycolytic pyruvate production. Interestingly, this manipulation did not rescue the loss of beta cell identity (Figure 4 k), prompting further investigation.

### Presenilin-1 controls the glycolytic flux and contributes to preservation of beta cell identity by regulating malate-aspartate shuttle activity

We and others have shown that a reduction in mitochondrial intermembrane space (IMS) Ca^2+^ level can inhibit the activity of IMS residing Ca^2+^ sensitive NADH shuttles, which in turn can influence glycolysis^4,5^. As we have observed several indications of disrupted Ca^2+^ homeostasis around mitochondria, we wondered if the IMS Ca^2+^ level is affected by PS1. PS1 KD resulted in reduced basal IMS Ca^2+^ (Figure S4 a), which opened a new inquiry into the mechanism of PS1 dependent control of beta cell glycolysis.

To test if reduced Ca^2+^ homeostasis at mitochondrial IMS could have impaired NADH shuttle activity, we measured the cytosolic NAD^+^/NADH ratio in INS-1 cells using genetically encoded biosensor Peredox^41^. PS1 KD resulted in a glucose-dependent reduction of cytosolic NAD^+^/NADH ratio (Figure 5 a), suggesting an inability of these cells to recycle cytosolic NADH back to NAD^+^. This metabolic defect may be the reason behind the hampered glycolytic flux caused by PS1 KD. Thus, we hypothesized that PS1 controls glycolysis by maintaining IMS Ca^2+^ homeostasis and supporting the activity of Ca^2+^-sensitive NADH shuttles in the mitochondrial IMS. These shuttles recycle cytosolic NADH back to NAD^+^, supplying NAD^+^ to maintain the activity of glyceraldehyde-3-phosphate dehydrogenase (GAPDH). Accordingly, we suspected a bottleneck within glycolysis in the GAPDH-catalyzed step in PS1 KD cells (Figure 5 b).

**Figure 5.**
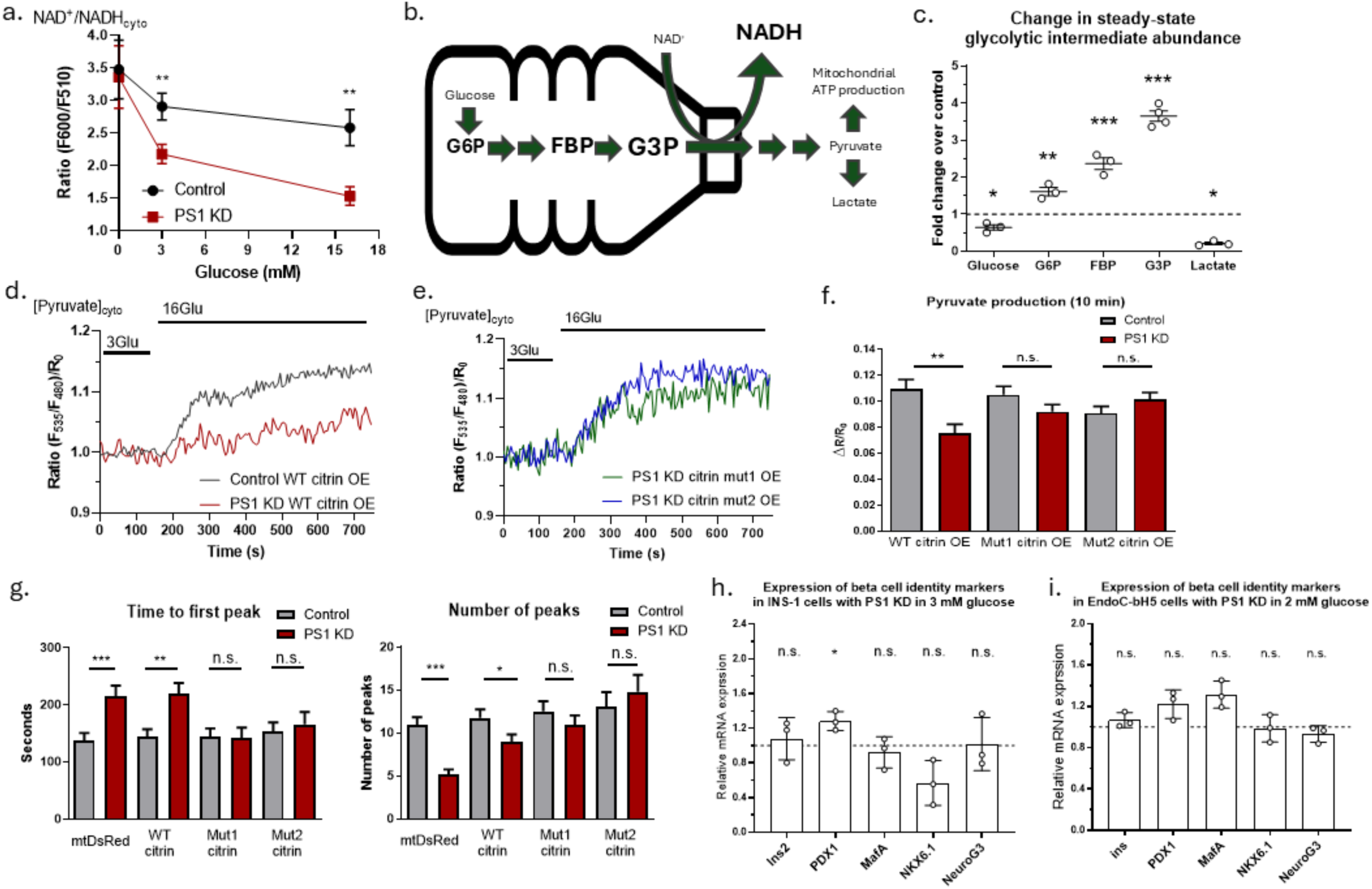
Presenilin-1 controls the glycolytic flux and contributes to preservation of beta cell identity by regulating malate-aspartate shuttle activity. **a,** glucose dependent changes in the cytosolic NAD^+^/NADH ratio shown as the Peredox biosensor emission ratio (+/-SEM) for control (black) and PS1 KD (red) INS-1 cells (control: 0 glucose n=8, 3 glucose n=11, 16 glucose n=10; PS1 KD: 0 glucose n=8, 3 glucose n=11, 16 glucose n=10). **b,** graphical representation of the hypothesized glycolytic bottleneck in PS1 KD beta cells; font size represents metabolite abundance. **c,** statistical analysis of mass spectrometry assay of glycolytic intermediates in INS-1 cells; metabolite abundance of PS1 KD cells is normalized to control (n=3-4). **d**, representative cytosolic pyruvate traces in response to increased extracellular glucose in control (black) and PS1 KD (red) INS-1 cells overexpressing wild type citrin construct. **e**, representative cytosolic pyruvate traces in response to increased extracellular glucose in PS1 KD INS-1 cells overexpressing wild mutant citrin constructs. **f**, statistical analysis of the experiments shown in **d** and **e**, bar graphs (+/-SEM) show changes in cytosolic pyruvate after 10 min in 16 mM glucose (control: WT citrin OE n=37, mut1 citrin OE n=38, mut2 citrin OE n=42; PS1 KD: WT citrin OE n=38, mut1 citrin OE n=45, mut2 citrin OE n=46). **g**, bar graphs (+/-SEM) on the left panel show time delay between the addition of 16 mM glucose and the onset of Ca^2+^ oscillations in INS-1 cells; bar graphs (+/-SEM) on the right panel show the number of Ca^2+^ oscillations/peaks in INS-1 cells (control: mtDsRed OE n=92, WT citrin OE n=79, mut1 citrin OE n=75, mut2 citrin OE n=53; PS1 KD: mtDsRed OE n=84, WT citrin OE n=74, mut1 citrin OE n=46, mut2 citrin OE n=41). Expression of beta cell identity markers in PS1 KD INS-1 cells in 3 mM glucose (**h**) and EndoC-bH5 cells in 2 mM glucose (**i**); values are normalized to respective average WT expression under the same experimental condition, n=3. Unpaired t-test (**a-g**), paired t-test (**h, i**), n.s. p > 0.05, * p < 0.05, ** p < 0.01, *** p < 0.001.

To test our hypothesis, we measured glycolytic intermediates using mass spectrometry. Our analysis revealed the accumulation of metabolites between glucose 6-phosphate (G6P) and glyceraldehyde 3-phosphate (G3P), and the reduction of lactate in PS1 KD (Figure 5 c), supporting our bottleneck hypothesis. Curiously, the analysis also showed decreased glucose level in PS1 KD cells. Since no defects in glucose uptake upon PS1 KD were found, we assumed that it could be an adaptive mechanism in response to hampered glycolytic flux. No glycolytic intermediates between G3P and lactate were detectable, likely due to low abundance, which can also explain the fact that we observed no significant difference in basal pyruvate level between control and PS1 KD cells in 3 mM glucose.

In order to test our proposed mechanism of PS1 mediated control of glycolysis in a more direct manner, we attempted to rescue the glycolysis by manipulating the malate-aspartate shuttle (MAS). MAS is the main NADH shuttle in pancreatic beta cells, which express aspartate-glutamate mitochondrial carrier 1 (AGC1, or aralar), as the Ca^2+^ sensitive component^42–44^. AGC1 is half-maximally stimulated at around 300 nM Ca^2+^ ^44^ and thus we wondered if we could rescue glycolysis in PS1 KD cells by overexpressing AGC2 (citrin), a homolog of AGC1 in non-excitable cells, which requires only around 100 nM Ca^2+^ for half-maximal stimulation^44^. Using pyruvate production upon glucose elevation in INS-1 cells as a readout, we found out that AGC2 overexpression (OE) did not restore pyruvate production of PS1 KD cells (Figure 5 d), suggesting that wild-type AGC2 might still be influenced by change in Ca^2+^ level in IMS. Consequently, we generated AGC2 mutants that can potentially mimic its conformational change upon Ca^2+^ binding. For that, we relied on available AGC1-2 crystal structures^45^ and introduced single amino acid exchanges at glutamine 67 (mutant 1) or glutamine 76 (mutant 2) to lysine. OE of either of these mutants in PS1 KD cells ultimately rescued the pyruvate production upon glucose stimulation (Figure 5 e, f, Figure S4 b), supporting our hypothesis on PS1-mediated control of glycolytic flux through modulation of Ca^2+^ sensitive MAS activity. Furthermore, OE of Ca^2+^ bound state mimicking AGC2 mutants, and not wild-type (WT) AGC2, rescued glucose-stimulated Ca^2+^ oscillations in PS1 KD INS-1 cells (Figure 5 g), further confirming the mechanism of PS1 mediated control of beta cell glucose metabolism.

Finally, to test if the glycolytic bottleneck could cause the loss of identity observed in PS1 KD cells, we reduced culture glucose concentration. Interestingly, significantly lowering culture glucose concentration rescued reduced expression of beta cell identity markers in INS-1 and EndoC-bH5 cells (Figure 5 h, i, Figure S4 c), pointing at the glycolytic bottleneck as being responsible for the loss of beta cell identity in beta cells with PS1 KD.

## Discussion

The role of the pancreas as a driver of bodily metabolism is dependent on its resident cells’ ability to detect and respond to certain metabolic cues. Pancreatic beta cells detect and respond to elevation in blood glucose level by releasing insulin, thus steering whole body glucose metabolism. Inability of the beta cells to properly respond to elevations of blood glucose can lead to drastic metabolic phenotypes, including dysfunction of the CNS^46,47^. Our previous work on the function of PS1 in pancreatic beta cells left open questions regarding the detailed mechanism of PS1-mediated control of mitochondrial bioenergetics as well as the involvement of PS1 in glucose responsiveness of beta cells^16,17^. To address these questions and to shed light into the relevance of AD causing mutations on beta cell function and identity, we utilized pancreatic islets and organ slices of an *in-vivo* knock-in mouse model of Alzheimer’s disease (3xTg AD), which carry the familial AD associated PS1 M146V mutation in the whole body, including the pancreas, along with mutated APP and Tau with a brain-specific expression profile^20^. For detailed mechanistic analysis, we used rat and human pancreatic beta cell lines, INS-1 and EndoC-bH5, respectively. Our previous work has demonstrated that INS-1 rat insulinoma cell line can be reliably used to replicate beta cell physiology and are comparable to freshly isolated pancreatic islets^16,17^. Additionally, key experiments were repeated using transient KD of PS1 by adenoviral delivery of shRNA in primary pancreatic islets and organ slices of WT mice. To decipher the intricate control mechanisms of beta cell metabolism and the role of PS1, we took advantage of recent advances in live-cell imaging using various ion and metabolite biosensors, where INS-1 cell line offered the most flexibility in terms of expression and robustness.

It has been reported that diabetes is a risk factor of dementia, including Alzheimer’s disease^11,14,46,47^. In fact, the two diseases share very common metabolic signatures^11^. Given that PS1 mutations are the major cause of familial AD^48^, studying the role of PS1 mutations in the pancreatic beta cells could give us a better understanding of the protein’s role in both organs and diseases. The triple transgenic mouse model of AD used in this study contains human PS1 M146V mutant knock-in protein^20,21^, reported to cause loss of ER Ca^2+^ leak function^15^, thus we suspected that it would mimic some of the PS1 KD phenotypes we have previously observed. In line with our experiments with transient KD of PS1 (Figure S1 c-e), islets from 3xTg FAD model mice resulted in delayed and reduced glucose-induced Ca^2+^ oscillations (Figure 1 a-c). A delay in the onset of Ca^2+^ oscillations can have substantial implications, such as delayed initial insulin release (Figure 1 d). Although we did not obverse significant changes in prolonged insulin release, insulin content, and blood glucose concentration in the 3xTg AD model, delayed or reduced initial phase of insulin release (Figure 1 d-f) could point to an onset of diabetes^49,50^. Indeed, previous research has demonstrated impaired glucose homeostasis and changes in fasting insulin in 3xTg AD model mice and one report also suggested the presence of human APP/amyloid beta in the pancreas of these mice^51,52^. Whether or not these potential APP/amyloid beta deposits contribute to beta cell metabolic changes in concert with presenilin mutation requires further research.

Since the loss of beta cell identity is a hallmark of diabetes^7,9^, we investigated if there is an indication of anonset of a loss of identity in the islets of AD model mice. Expression analysis of beta cell identity genes revealed a trend pointing towards a loss of beta cell identity in the pancreas of 3xTg mice (Figure S1 a), in line with previous observations of reduced pancreatic insulin expression^52^. We assumed that the transgenic knock-in models might have developed compensatory mechanisms to preserve their beta cell identity. To address this possibility, we analyzed beta cell identity changes in primary mouse islets and rat and human cell beta cell lines with transient KD of PS1. The experiments revealed that indeed PS1 was necessary to preserve identity of pancreatic beta cells (Figure 1 g-j). These results prompted us to start a detailed analysis of the role of PS1 in preservation of beta cell identity.

In pursuit of the mechanism behind the loss of beta cell identity, we searched for known pathways that influence beta cell identity and could be linked to Ca^2+^ leak function of PS1. Using available biosensors monitoring/reporting the activity of CREB and CamKII, we found that PS1 KD did not affect them (Figure S2 a-d). Curiously, nuclear translocation of NFATc2 and c3 was increased (Figure 2 a, b), which was surprising for two reasons: first, PS1 KD leads to reduced ER Ca^2+^ leak and glucose stimulated Ca^2+^ signaling, and second, NFAT signaling was reported to preserve beta cell identity^53^. This result made us re-evaluate sub-cellular Ca^2+^ signatures in beta cells with PS1 KD, as it could reveal the reason for increased NFAT activation. Using Ca^2+^ biosensors with various sensitivities and targeting techniques, we deciphered an altered sub-cellular Ca^2+^ oscillation pattern, where sub-PM region of PS1 KD cells had increased Ca^2+^ oscillations of small amplitude (Figure 2 d), which were the reason for calcineurin and NFAT activation (Figure 3 a).

While the calcineurin-NFAT pathway was shown to positively affect the beta cell development and identity^53^, calcineurin has many substrates. Among them is AMPK, which can be dephosphorylated by calcineurin, thereby reducing its activity^27,28^. The fact that AMPK activity was lowered by PS1 KD despite reduced cellular ATP^16^ suggests that inhibition of AMPK by calcineurin exceeds its activation by reduced cellular energy status (Figure 3 b). Several recent publications highlight the importance of AMPK in beta cell development and maintenance of beta cell identity^30–32,54^. Additionally, the balance between AMPK and mTORC1 pathways plays an important role during beta cell development, where AMPK suppresses mTORC1 activity post-weaning and into maturation of beta cells^32,33^. Thus, the fact that our attempt at normalization of the AMPK and mTORC1 pathways did not rescue beta cell identity (Figure 3 d-g) suggests that an alternative mechanism of PS1 dependent control has a stronger impact on beta cell identity.

Metabolism is a key factor in cell fate determination^55,56^. In pancreatic beta cells, accumulation of glycolytic intermediates between fructose-bisphosphate and glyceraldehyde-triphosphate as a result of hyperglycemia was shown to initiate beta cell dedifferentiation by activating mTORC1^34^. Our work shows that the same glycolytic intermediates can accumulate as a result of a bottleneck at the GAPDH catalyzed step, despite reduction in the total glycolytic flux. We attempted to metabolically reverse the effect of PS1 KD on glycolytic flux, either by supplying extra pyruvate, which should rescue Ca^2+^ signaling (Figure 4 a, b), or by reducing glucose concentration, which should reduce the accumulation of glycolytic intermediates. Of the two metabolic manipulations, lowering glucose concentration from 10 mM to 3 mM rescued the identity loss in PS1 KD cells (Figure 5 h, I, Figure S4 c). These results suggest that PS1 contributes to the maintenance of beta cell identity via controlling their glycolytic flux by regulating the activity of malate-aspartate shuttle (Figure 5 b-g).

In summary, the current study identifies novel roles of PS1 in pancreatic beta cells, which include control of glucose metabolism through regulation of malate-aspartate shuttle activity and regulation of various Ca^2+^ dependent signaling pathways. Furthermore, we found that PS1 exerts pressure on beta cell identity preservation via controlling the glycolytic flux. Thus, the current work provides a new perspective into the role of PS1, a risk factor of AD, in systemic glucose metabolism, opening potential research and therapeutical avenues in AD field.

## SUPPLEMENTAL DATA

**Supplementary figure 1.**
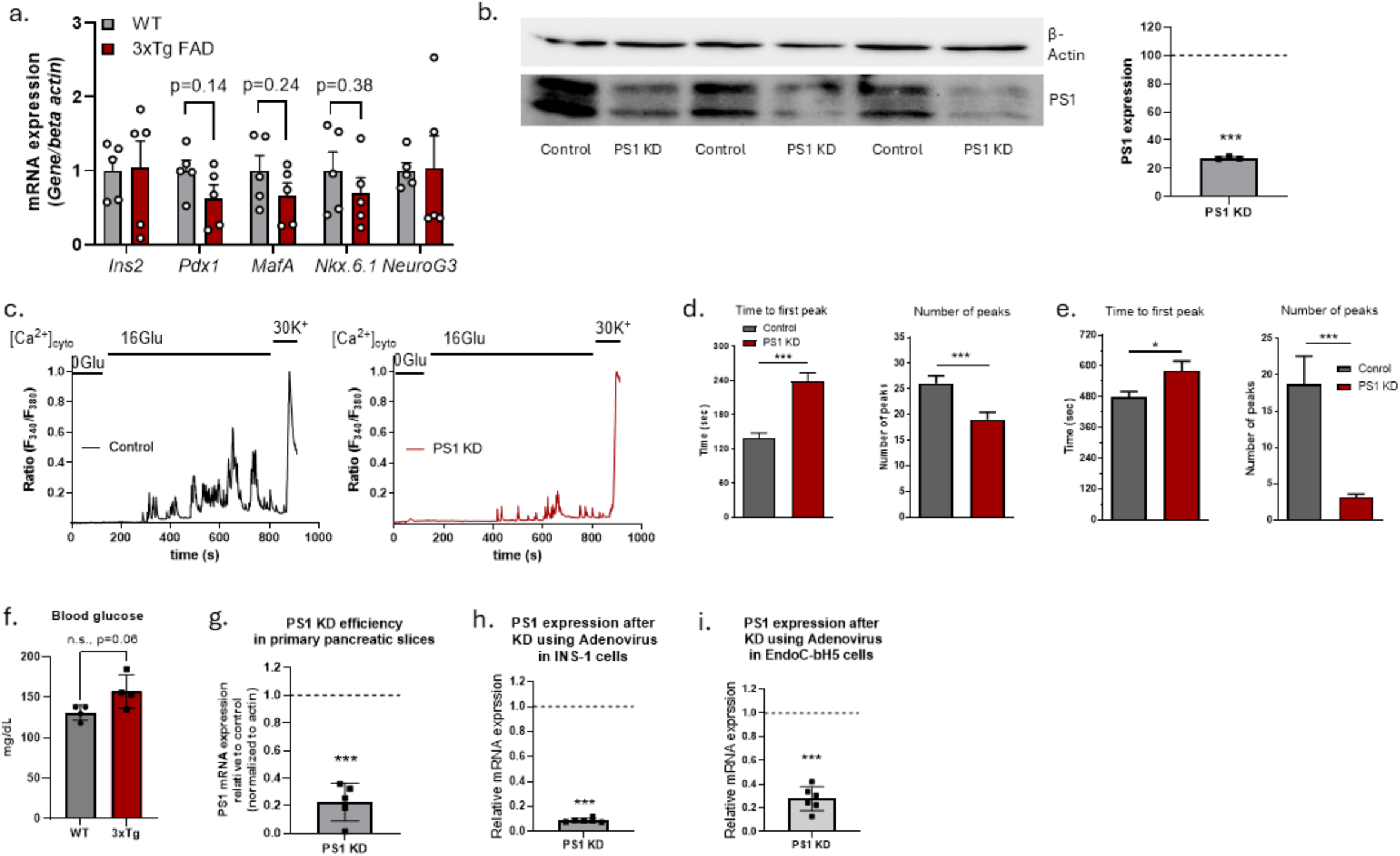
**a**, expression of beta cell identity markers in pancreatic islets from WT and 3xTg FAD; values are normalized to respective average WT expression (n=5). **b**, Western blot (left) and analysis (right) show the PS1 expression in control and PS1 KD INS-1 cells (n=3). **c**, representative FURA-2 AM traces depicting cytosolic Ca^2+^ oscillations in response to 16 mM glucose in control (left) and PS1 KD (right) INS-1 cells. **d**, statistical analysis of experiments as shown in **c,** bar graphs (+/-SEM) on the left panel show time delay between the addition of 16 mM glucose and the onset of Ca^2+^ oscillations; bar graphs (+/-SEM) on the right panel show the number of Ca^2+^ oscillations/peaks (control, n=157, PS1 KD, n=167). **e**, statistical analysis of dynamic Ca^2+^ measurements done with primary pancreatic slices; bar graphs (+/-SEM) on the left panel show time delay between the switch from 6 mM to 8 mM glucose and the onset of Ca^2+^ oscillations; bar graphs (+/-SEM) on the right panel show the number of Ca^2+^ oscillations/peaks (control, n=164, PS1 KD, n=148). **f**, Blood glucose level in WT and 3xTg FAD mice, n=4. PS1 mRNA KD efficiency in primary mouse pancreatic organ slices (**g**, n=5), INS-1 cells (**h**, n=6), and EndoC-bH5 cells (**i**, n=6). Unpaired t-test (for **d-f**), paired t-test (for **b, g-i**), n.s. p > 0.05, * p < 0.05, ** p < 0.01, *** p < 0.001

**Supplementary figure 2.**
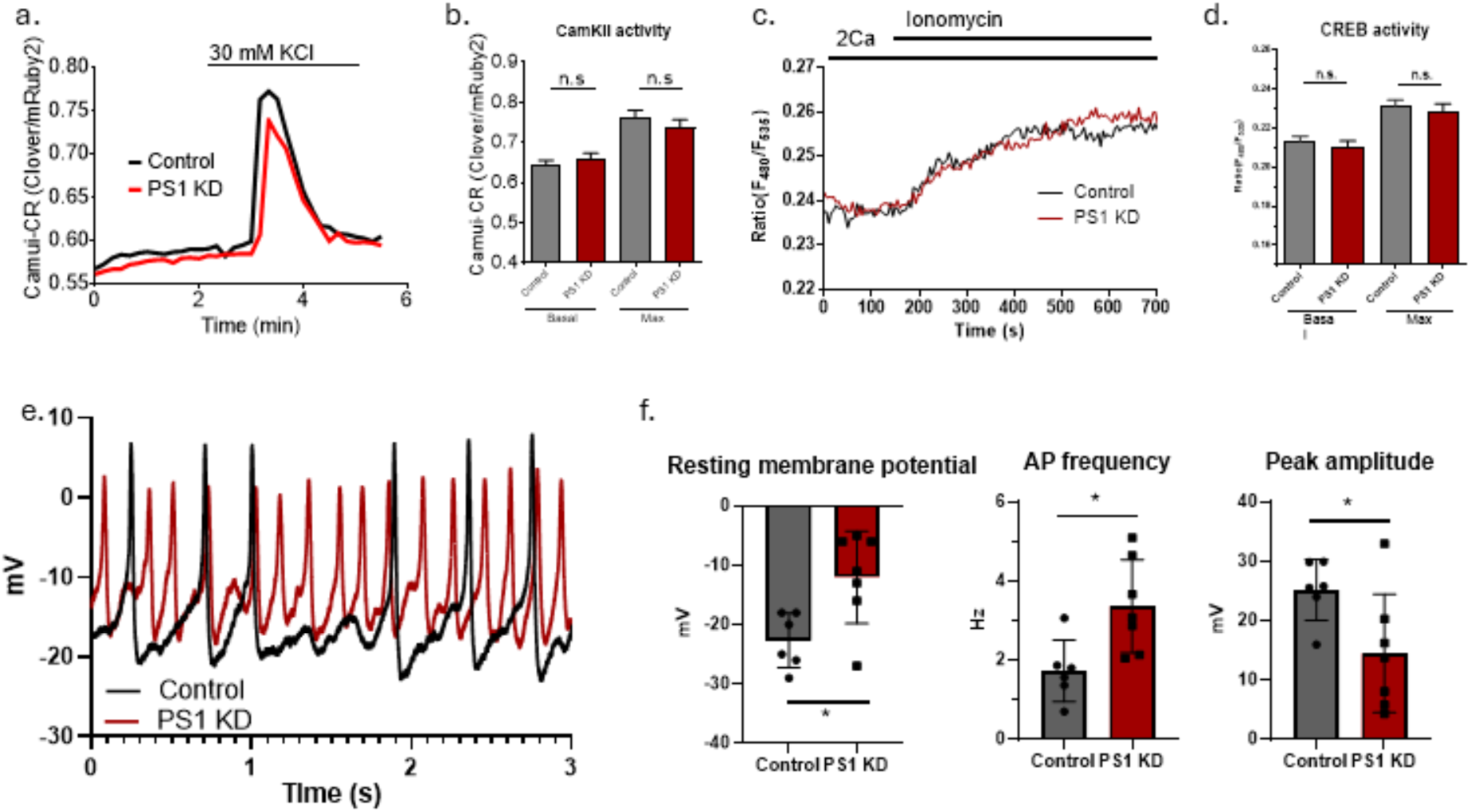
**a,** representative traces depicting basal and stimulated (with 30 mM KCl) activity of CamKII, measured with CamKII activity biosensor Camui-CR^68^. **b,** statistical analysis of basal and stimulated CamKII activity in control and PS1 KD INS-1 cells (control n=45, PS1 KD n=47). **c**, representative traces depicting basal and stimulated (with 4 uM ionomycin) activity of CREB, measured with CREB activity biosensor NLS-ICAP^67^. **d,** statistical analysis of basal and stimulated CREB activity in control and PS1 KD INS-1 cells (control n=36, PS1 KD n=37). **e,** representative membrane potential traces of INS-1 cells. **f,** statistical analysis of resting membrane potential, AP frequency and amplitude in control and PS1 KD INS-1 cells (n=6). (control n=26, PS1 KD n=27). Unpaired t-test, n.s. p>0.05, **p < 0.01.

**Supplementary figure 3.**
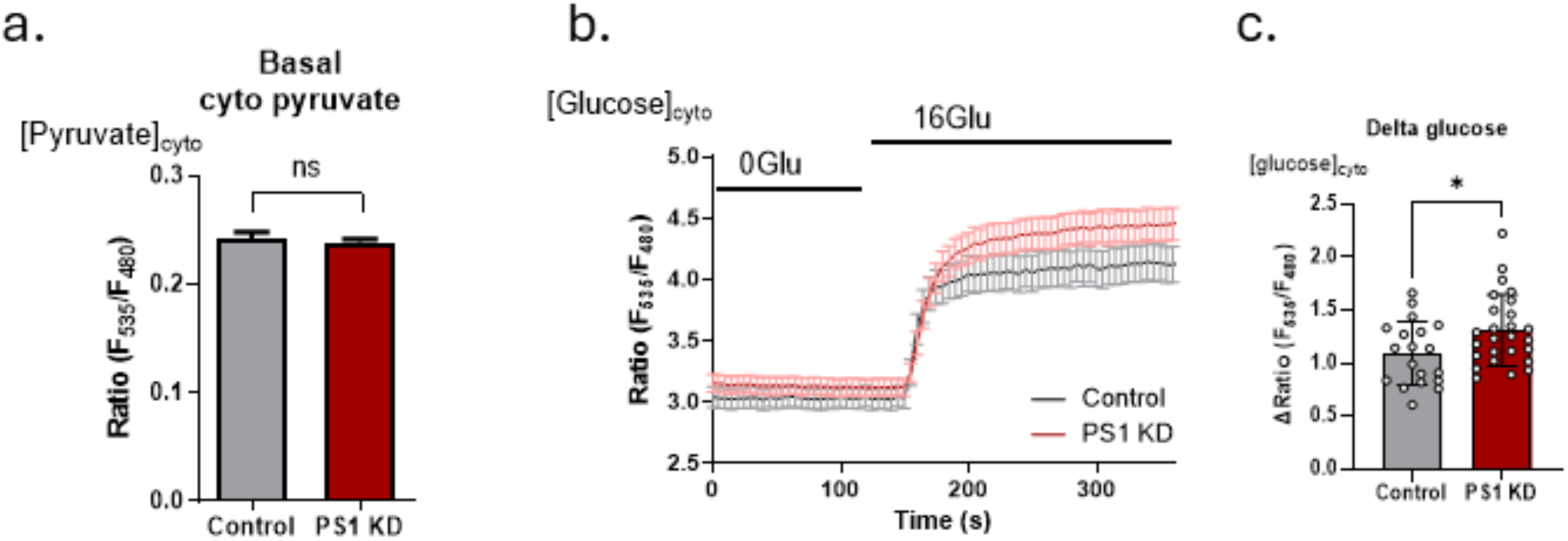
**a,** basal cytosolic pyruvate levels shown as Pyronic emission ratio in control and PS1 KD INS-1 cells (control, n=43; PS1 KD, n=71). **b**, cytosolic glucose dynamics (mean +/-SEM) shown as FLII12Pglu-700μδ6 emission ratio upon extracellular glucose addition in control and PS1 KD INS-1 cells. **c**, change in cytosolic glucose level (+/-SEM) after addition of 16 mM extracellular glucose as shown in **b** (control, n=19; PS1 KD, n=26). Unpaired t-test, n.s. p>0.05, *p <0.05.

**Supplementary figure 4.**
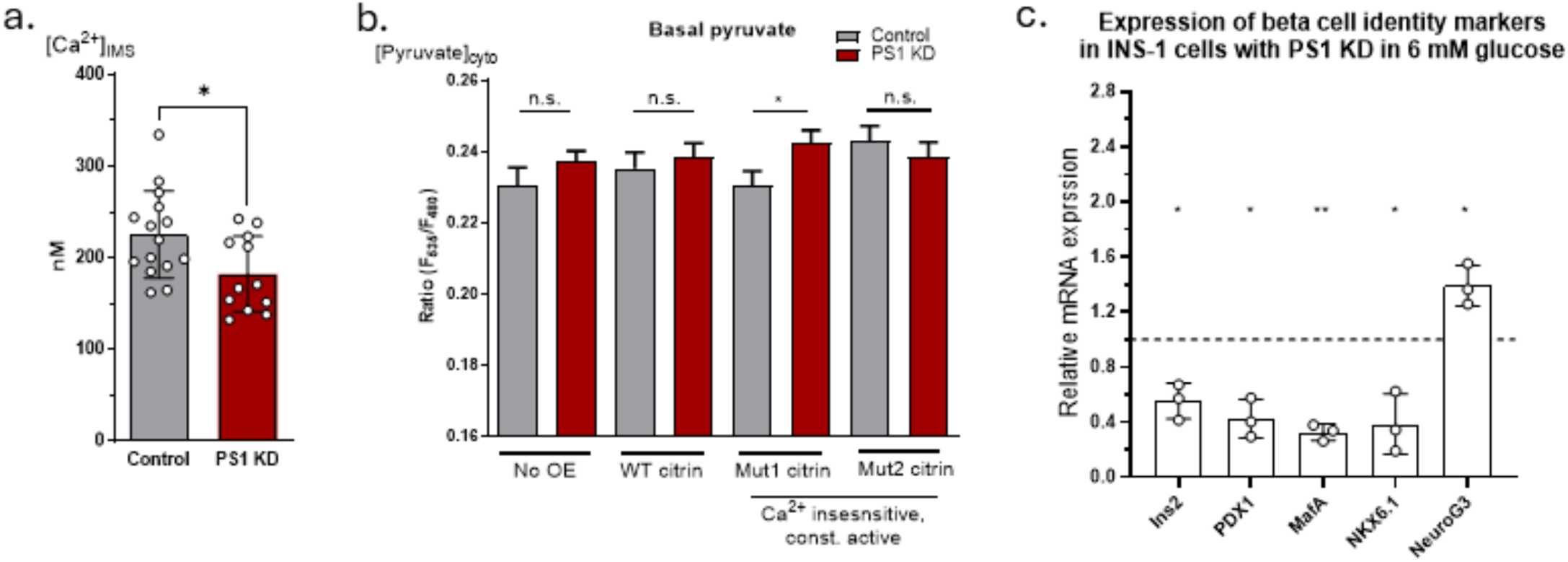
**a,** bar graphs (+/-SEM) depict basal Ca^2+^ concentrations in the mitochondrial IMS (control, n=15, PS1 KD, n=12) of INS-1 cells. **b**, basal cytosolic pyruvate levels shown as Pyronic emission ratio in control and PS1 KD INS-1 cells with OE of WT and mutant citrin constructs (control: no OE n=67, WT citrin OE n=38, mut1 OE n=38, mut2 OE n=42; PS1 KD: no OE n=88, WT citrin OE n=38, mut1 OE n=45, mut2 OE n=47). **c**, expression of beta cell identity markers in PS1 KD INS-1 cells in 6 mM glucose; values are normalized to respective average WT expression under the same experimental condition, n=3. Unpaired t-test (**a** and **b**), paired t-test (**c**), n.s. p > 0.05, * p < 0.05, ** p < 0.01)

## Materials and methods

### Animals

All protocols were approved by the Federal Ministry of Education, Science and Research of the Republic of Austria (permit number: 2020-0.258.669). C57BL/6J mice (034830-JAX, Jackson Laboratories) and 3xTg-AD mice (034830-JAX, Jackson Laboratories) were used to prepare pancreatic tissue slices. All animals were kept on a 12:12-h light:dark schedule in individually ventilated cages (Allentown LLC).

Breeding pairs of triple transgenic mice (3xTg), carrying the PS1M146V, APPSwe, and tauP301L transgenes (Oddo et al, 2003), were initially purchased from The Jackson Laboratories (JAX), United States of America. Transgenic animals used in this study were bred in the animal facility of the Institute of Molecular Biosciences, Graz, Austria, using the breeding pairs from Charles River Laboratories. The control strain used for the AD disease model was B6129SF2/J, also bred in-house with breeding pairs from Charles River Laboratories. Animals were housed under conditions essentially as described^57^ and all animal experiments were performed in accordance with national and European ethical regulations (Directive 2010/63/EU) and approved by the responsible institutional or government agencies (Bundesministerium fur Wissenschaft, Forschung und Wirtschaft, BMWFW, Austria: BMWFW-66.007/0032-V/3b/2019, GZ 2023-0.483.703 and GZ 2023-0.691.200).

### Cell Culture and Transfection

The INS-1 832/13 (INS-1) cells were a generous gift from Prof. Dr. Claes B. Wollheim and Dr. Françoise Assimacopoulos-Jeannet (University Medical Center, Geneva, Switzerland). INS-1 cells were cultured in RPMI 1640 containing 10 mM glucose (PubChem CID: 5793) supplemented with 10 mM HEPES (PubChem CID: 23831), 10% fetal calf serum (FCS), 1 mM sodium pyruvate (PubChem CID: 23662274), 50 μM β-mercaptoethanol (PubChem CID: 1567), 1% (v/v) Pen Strep® (ThermoFischer, Vienna, Austria; 10.000 U/L), and 1.25 μg/ml Amphotericin B (ThermoFischer, Vienna, Austria; 250 μg/mL). Cells were used between passage numbers 53 and 68. Human EndoC-bH5 cells were purchased from Human Cell Design, Inc., and cultured according to the manufacturer’s guidelines in culture media with 5.5. mM glucose. All cells were kept in a humidified incubator (37°C, 5% CO_2_, 95% air).

For all microscopic experiments, INS-1 cells were plated on 30 mm glass coverslips in 6-well plates and transfected at 50–60% confluency with biosensor DNA constructs (1 μg/well) alone or with siRNA against PS1/scramble control (Table 1) by using 3 μl TransFast transfection reagent (Promega, Madison, WI, USA) in 1 mL of serum and antibiotic-free medium for 12–14 h. After that, transfection media was replaced with 2 mL of full culture medium. All experiments were performed 40-45 h after transfection. Human EndoC-bH5 cells were transduced with adenovirus carrying specific PS1 shRNA (Table 1) or scrambled shRNA (VectorBuilder) and cultured for 48 h. Custom adenovirus constructs were generated that carry a reporter (GFP or mCherry) or a genetically encoded biosensor PyronicSF along with scrambled shRNA or shRNA against PS1 along. These adenoviruses were used for EndoC-bH5 and primary mouse pancreatic slices.

**Table 1.**
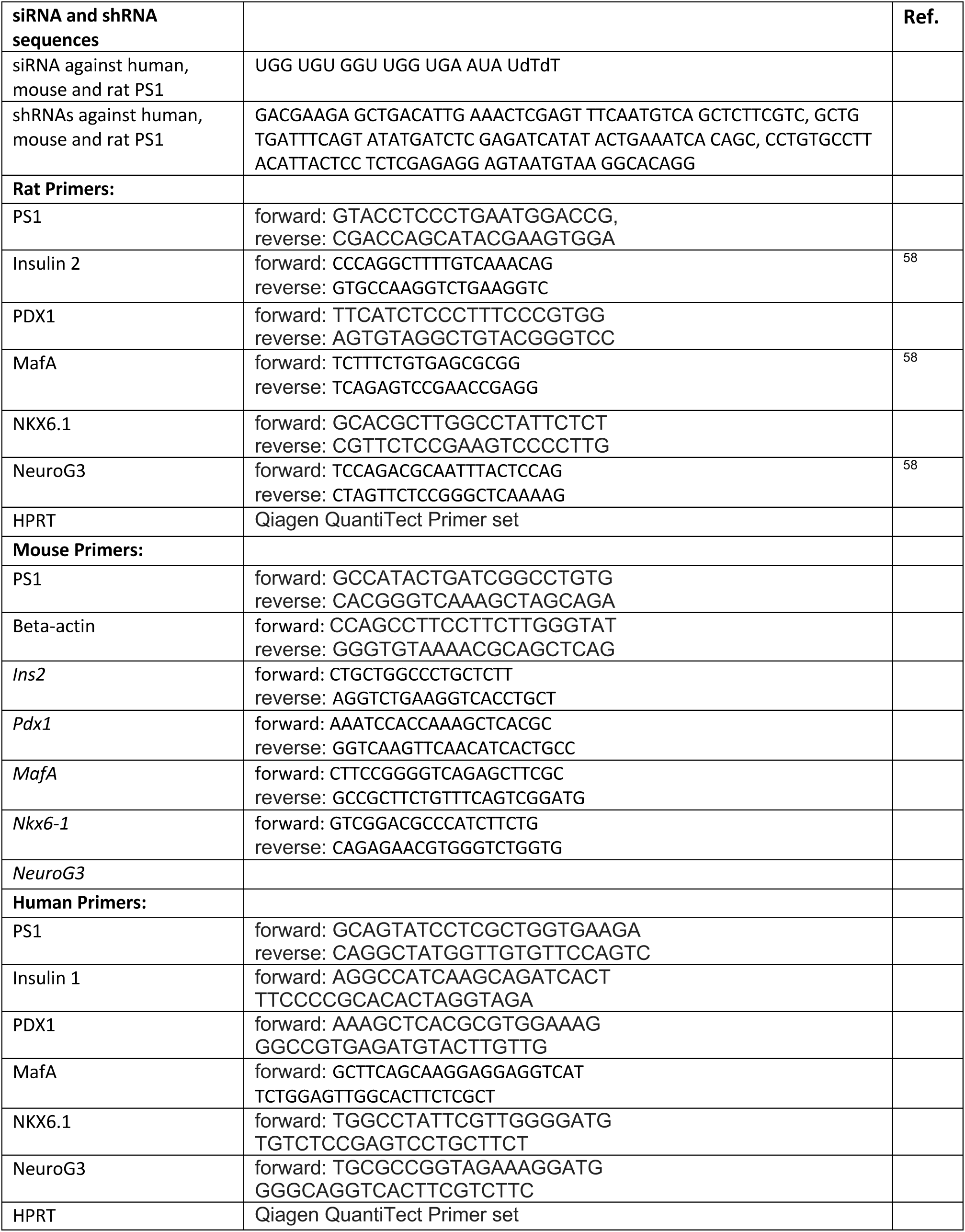

### Live cell imaging of INS-1 and EndoC-bH5 cells

All live-cell microscopy experiments were performed on an Olympus IX73 inverted microscope if not mentioned otherwise. The microscope is equipped with an UApoN340 40× oil immersion objective (Olympus, Japan) and a CCD Retiga R1 camera (Q-imaging, Canada). For illumination, a LedHUB® (Omnicron, Germany) equipped with 340, 385, 455, 470, and 550 nm LEDs in combination with CFP/YFP/RFP (CFP/YFP/mCherry-3X, Semrock, USA) or GFP (GFP-3035D, Semrock, USA) filter set was used. During the measurements cells were continuously perfused by a gravity-based perfusion system (NGFI, Graz, Austria). Data acquisition and control of the fluorescence microscope was performed using Visiview 4.2.01 (Visitron, Germany). Background subtracted regions of interest were used for analysis. During live cell imaging experiments, cells were continuously perfused with physiological buffer containing 2 mM CaCl, 135 mM NaCl, 1 mM MgCl_2_, 5 mM KCl, 10 mM HEPES, and indicated glucose concentrations with pH adjusted to 7.4 at room temperature.

Basal IMS Ca^2+^ levels were measured and quantified as previously reported by us^4^ using IMS targeted GEM-GECO^58^ biosensor. Absolute Ca^2+^ concentrations were quantified by obtaining minimum and maximum ratio changes of the biosensor in the presence of ionomycin with 0 and 2 mM Ca^2+^, respectively (0 Ca^2+^ buffer contained 100 µM EGTA).

Cytosolic Ca^2+^ oscillations in response to high glucose and pyruvate were measured using FURA-2 AM Ca^2+^ sensitive dye as previously reported^17^. Shortly, FURA2-AM loading was done in a storage buffer (2 mM CaCl, 138 mM NaCl, 1 mM MgCl_2_, 5 mM KCl, 10 mM HEPES, 2.6 mM NaHCO3, 0.44 mM KH_2_PO_4_, amino acid, and vitamin mix, 3 mM glucose, 2 mM L-glutamine, 1% Penicillin/Streptomycin, 1% Fungizone, pH adjusted to 7.4) with 3 µM FURA2-AM for 30 min at room temperature and washed with fresh storage buffer. FURA2-AM was sequentially excited with 340 nm and 385 nm LEDs and emission collected with GFP emission filter set.

Basal pyruvate and pyruvate production in response to 16 mM glucose in INS-1 cells was quantified using genetically encoded pyruvate biosensor Pyronic^36^. Pyruvate production in EndoC-bH5 cells, pyruvate flux in INS-1 cells, and pyruvate flux in primary mouse pancreatic slices was quantified using PyronicSF biosensor^37^. Adenoviral delivery of biosensors and shRNA was used for human EndoC-bH5 cells and primary mouse pancreatic slices. Cells expressing Pyronic were excited with 455 nm LED and emission collected at 480 nm and 530 nm using a CFP/YFP/RFP filter set and 505dcxr beam-splitter. PyronicSF was excited with 470 nm LED and emission collected with a GFP filter set.

Cytosolic NAD^+^/NADH ratio was quantified in INS-1 cells at given glucose concentrations using NAD^+^/NADH ratio biosensor Peredox^41^ as previously described^4^. INS-1 cells expressing Peredox were excited with 385 nm and 550 nm LEDs and emission was collected using a CFP/YFP/RFP filter set and 565 LPXR beam-splitter (Semrock, USA).

Nuclear translocation of NFAT in INS-1 cells under resting conditions was done on confocal spinning disk microscope (Axio Observer.Z1 from Zeiss, Gottingen, Germany) while the cells were in kept in storage buffer. The confocal microscope is equipped with a 100x objective lens (Plan-Fluor x100/1.45 Oil, Zeiss), a motorized filter wheel (CSUX1FW, Yokogawa Electric Corporation, Tokyo, Japan) on the emission side, AOTF-based laser merge module for laser line 405, 445, 473, 488, 561, and 561 nm (Visitron Systems) and a Nipkow-based confocal scanning unit (CSU-X1, Yokogawa Electric Corporation). A ratio of nucleus ROI to cytosolic ROI was used to analyze the nuclear translocation ratio of GFP-tagged NFAT constructs.

Calcineurin activation was measured using genetically encoded calcineurin activation status biosensor CaNARi, targeted to OMM, cytosol, and subPM^26^. The sensor was excited with 455 nm LED and emission was collected at 480 nm and 530 nm using a CFP/YFP/RFP filter set and 505dcxr beam-splitter. Cells were imaged in the storage buffer.

AMPK activity was measured using genetically encoded AMPK activity biosensor AMPKAR^59^. The sensor was excited with 455 nm LED and emission was collected at 480 nm and 530 nm using a CFP/YFP/RFP filter set and 505dcxr beam-splitter. Cells were imaged in the storage buffer.

mTORC1 activity was measured using genetically encoded mTORC1 activity biosensor TORCAR^60^. The sensor was excited with 455 nm LED and emission was collected at 480 nm and 530 nm using a CFP/YFP/RFP filter set and 505dcxr beam-splitter. Cells were imaged in the storage buffer.

CREB activity was measured using genetically encoded nuclear targeted CREB activity biosensor NLS-ICAP^61^. The sensor was excited with 455 nm LED and emission collected at 480 nm and 530 nm using a CFP/YFP/RFP filter set and 505dcxr beam-splitter. Cells were imaged in the storage buffer.

CamKII activity was measured using genetically encoded CamKII activity biosensor Camui-CR^62^. The sensor was excited with 470 nm LED and emission was collected at 510 nm and 570 nm using a GFP/RFP filter set and 530 nm beam-splitter on AnglerFish microscope (NGFI, Graz, Austria).

### Pancreatic Tissue Slice Preparation

Following CO_2_ euthanasia, the mouse abdominal cavity was assessed via laparotomy. 1.9% low melting-point agarose (Lonza, USA) dissolved in extracellular solution (ECS, (in mM): 125 28 NaCl, NaHCO_3_, 6 glucose, 6 lactic acid, 3 myo-inositol, 2.5 KCl, 2 CaCl_2_, 1.25 NaH_2_PO_4_, 1 MgCl_2_, 0.25 ascorbic acid) was infused at 40°C into the pancreas via the distally clamped common bile duct. The pancreas was then cooled, excised, and sliced using a vibratome (VT1000S, Leica Biosystems). The sliced pancreas (140 µm) was maintained in HEPES-ECS (in mM: 125 NaCl, 10 NaHCO_3_, 10 HEPES, 6 glucose, 6 lactic acid, 3 myo-inositol, 2.5 KCl, 2 Na-pyruvate, 2 CaCl_2_, 1.25 NaH_2_PO_4_, 1 MgCl_2_, 0.25 ascorbic acid; titrated to pH 7.4 with 1 M NaOH). All chemicals were obtained from Sigma-Aldrich if not specified differently.

### Generation of PSEN1 Knockdown in Pancreatic Tissue Slices

Pancreatic tissue slices were cultured in RPMI medium without glucose (11879020, Thermo Fisher Scientific) and supplemented with 10% FBS (F7524, Sigma-Aldrich), 1% Penicillin/Streptomycin (15140122, Gibco, 1:100) and 6 mM glucose (G8270, Sigma-Aldrich) for 24-48h at 37°C.

To generate a PS1 knockdown in pancreatic tissue slices we transduced the slices with 5×10^6^ PFU/ml adenovirus carrying specific mouse PS1 shRNA (Ad-GFP-U6-m-PSEN1-shRNA, shADV-269593, Vector Biolabs) or scrambled shRNA (Ad-U6-RNAi-GFP,1122N, Vector Biolabs) and cultured for 24-48h.

### Dynamic [Ca^2+^]c Imaging in Mouse Pancreatic Tissue Slices

Pancreatic tissue slices were loaded with fluorescent Ca^2+^ indicator in HEPES-ECS (6.35 µM Calbryte 520 or 590-AM (AAT Bioquest), 0.03% Pluronic F-127 (w/v), and 36 0.12% dimethylsulphoxide (v/v)).

Dynamic [Ca^2+^]_c_ imaging was performed on a Nikon inverted confocal system (20×, NA 0.95). Each tissue slice was transferred into the imaging chamber that was continuously perfused with warm (34°C) HEPES-buffered extracellular solution supplemented with various glucose and pyruvate concentrations. The acquisition frequency was 30 Hz at 256×256 pixels resolution and approximately 1 µm^2^ pixel size. Fluorescence was excited with a 488 nm laser and detected at 500–700 nm for the Calbryte 520 indicator, or excited with a 561 nm laser and detected at 570–620 nm for the Calbryte 590 indicator, using a GaAsP PMT detector (Nikon A1R). Each islet [Ca^2+^]c dynamic recording lasted ∼3000s (∼50 minutes).

The analysis was performed as described in Postic at all, in brief, the regions of interest (ROIs) were semi-automatically determined by employing custom-made Python scripts (Python Software Foundation, Wilmington, Delaware, USA). From the selected ROIs we gather information about the [Ca^2+^]_c_ changes as the light intensity, the spatial coordinates of the ROIs and neighboring ROIs, statistic about the time recording, recording frequency, and pixel size.

Therefore, all the significant changes of [Ca^2+^]_c_ (with z-score>3) at all realistic time scales (limited by frequency and the time of the recording) within each ROI were automatically distilled and annotated as events. The latency to the first glucose-induced event was measured and compared across conditions. Event frequency was quantified as the number of events per minute per ROI.

### Dynamic pyruvate imaging in Mouse Pancreatic Tissue Slices

Pancreatic tissue slices were transduced with adenovirus containing scrambled or PS1 shRNA (described above) along with pyronicSF, intensiometric pyruvate biosensor. The slices were imaged after 24 h on a Leica TCS inverted confocal system (20×, NA 0.75). As before each tissue slice was transferred into the imaging chamber that was continuously perfused with warm (34°C) HEPES-buffered extracellular solution without lactate and pyruvate supplemented with glucose 6-8 mM and pyruvate transfer inhibitors UK 5099 and AR-C155858 (4186 and 4960, Tocris). The acquisition frequency was 0.2 Hz at 256×256 pixels resolution and approximately 1 µm2 pixel size. PyronicSF was excited by 490 nm line of a white laser and emitted light was detected by Leica HyD hybrid detector in the range of 500-700 nm using a photon counting mode (Leica Microsystems, Germany).

The recordings of pyruvate sensor in pancreatic organ slices were analyzed by ImageJ. First, we corrected for the bleaching, followed by hand-picking the ROIs containing the islet. The mean intensity was measured from the selected ROIs and used for further analysis.

### Quantitative PCR in INS-1 and EndoC-bH5 cells

Total mRNA was isolated using RNeasy® Mini Kit (Qiagen, Hilden, Germany), and reverse transcription was done using Applied Biosystems High-Capacity cDNA Reverse Transcription kit (Thermo Fisher Scientific Baltics UAB, Vilnus, Lithuania). qPCR was performed using Promega GOTaq® qPCR Master Mix (Madison, WI, USA). Primers used for quantification of expression of specific genes are listed in Table 1. HPRT1 was used for normalization. Primers were designed using NCBI primer design tool if not referenced otherwise.

### Quantitative PCR in primary pancreatic islets

RNA was transcribed into cDNA using Luna script RT supermix (NEB) according to the Manufacturer’s instructions. For qPCR analysis, 4 µl cDNA were mixed with 1 µl of respective forward and reverse primers, 6 µl Fresenius H_2_O, and 8 µl SYBR green (Bio-Rad) Target gene expression was calculated by the ΔΔCT method. β-actin was used as housekeeping gene.

### Western blot

For WB analysis, cells were seeded on 10 cm dishes, transfected with respective siRNAs and harvested 48 h post transfection. Primary antibodies were used at 1:1000 dilution: rabbit mAb Presenilin1 (D39D1, CST), mouse mAb b-Actin (sc-81178). Secondary antibodies were used at 1:5000 dilution: goat-anti-rabbit (sc-2054), m-IgGk BP-HRP (sc-516102). Broad Range (10-250 kDa) Color Prestained Protein Standard ladder (NEB, P7719S) was used in all blots.

### RT-PCR from the Mouse Pancreatic Tissue Slice

Mouse pancreatic tissue slices were examined and the ones containing were incubated as described. Following the incubation, we isolated the total RNA from pancreatic tissue slices (20 slices per condition) with Total RNA Kit, peqGOLD according to the manufacturer’s instruction (13-6834-02P, VWR). RNA was measured and adjusted to 500 ng/ml which was utilized in the reverse transcription reaction (4000 units, 10338842, INV) alongside dNTPs (100 mM each, R0181, Thermo Fisher) and random hexamers (50 µM, N8080127, Invitrogen). RT-PCR was performed for PS1 (fw: GCCATACTGATCGGCCTGTG; rv: CACGGGTCAAAGCTAGCAGA) using KAPA SYBR(R) FAST (KK4600, SIG) on the CFX Connect^TM^Real-Time System (Bio-Rad) and the following cycling parameters 20 sec 95°C, 20 sec 55°C and 1 sec 72°C for 40 cycles.

Samples were corrected for the total RNA input by normalizing CT values to the CT value of the β-actin (fw: CCAGCCTTCCTTCTTGGGTAT; rv: GGGTGTAAAACGCAGCTCAG).

### Immunofluorescence imaging

For blocking, the pancreatic tissue slices were incubated with blocking buffer (PBS containing 5% BSA (A2153, Sigma Aldrich)) supplemented with 0.1% Triton X-100 (37240, Serva Electrophoresis GmbH) for 1 hour at room temperature. After removal of the blocking solution, primary antibody was diluted in blocking buffer and added in the following concentrations: guinea-pig anti-insulin (ab7842, abcam, 1:100), rabbit anti-glucagon (A14609, ABclonal, 1:100), mouse anti-neurogenin (sc-374442, Santa Cruz Biotechnology 1:200), rabbit anti-PDX1 (5679S, Cell Signaling, 1:200), rabbit anti-MAFA (79737S, Cell Signaling, 1:500), rabbit anti-NKX6.1 (Cell Signaling, 54551S, 1:400). The slides were incubated with the primary antibody overnight at 4°C. The Secondary antibody incubation lasted for 1h at RT. Secondary antibodies were diluted in blocking buffer as follows: goat anti guinea-pig Alexa Flour 488 (A-11073, Invitrogen, 1:500), goat anti-rabbit Alexa Flour 647 (A-21244, Invitrogen, 1:500), goat anti-mouse DyLight 550 (SA510173, Sigma-Aldrich, 1:500). To label the nuclei, slices were incubated with 0.1 µg/ml DAPI (6335.1, Carl Roth). Images were obtained on a confocal microscope (Nikon A1R), using the excitation lasers at 405 nm, 488 nm, 561 nm and 640 nm and the GaAsP PMT detectors at 425–475 nm (DAPI), 500-550 nm (Alexa 488) 570–620 nm (DyLight 550) and 663-738 nm (Alexa 647). The possible crosstalk between the channels was minimized by sequential scanning. The slices were imaged with the z-stack mode, reaching about 30 µm in the tissue.

Each of the acquired optical planes was segmented into the ROIs representing single cells based on the DAPI staining with a custom-made ImageJ macro. The macro was developed in ImageJ for more information visit github.com/BHochreiter.

For each physical slice in a z-stack, we then choose the most representative optical plane: for each channel, we construct an average cumulative density function (CDF) of the log ROI intensities, as the median across optical planes. We then calculate the Kolmogorov-Smirnov (KS) distance of each of the optical planes, for each of the channels, as the maximal difference of the CDF to the average one. We sum the KS distances over channels and minimize over optical planes to identify the one, which has the CDF closest to the average CDF in all the channels. In most cases, this happens to be the middle optical plane.

For each channel, we then attempt to separate the background and signal ROIs: First, we infer the background intensity, by fitting the distribution of log ROI intensities to a two-component Gaussian, and choosing the lower mean value as the central background value to which we normalize all the values. This reduces to a simple subtraction in the log-intensity space. Finally, we use the fitted mixture of Gaussians to binarize the ROIs into background and signal, based on its intensity.

### Mass spectrometry

INS-1 cells were harvested and lysed in a 40:40:20 (V:V:V) mix of acetonitrile:methanol:water, extract was directly used for measurement. LC-MS Data was acquired on a Shimadzu LCMS-8060 system employing a ZICpHILIC (Merck) column (2.1 x 150 mm, 5 µm particle size). The flow rate was kept constant at 200 µL/min, Solvent A was H_2_O, 20 mM NH_4_(H)CO_3_, pH = 9.4, solvent B was acetonitrile, 20 mM NH_4_(H)CO_3_, pH = 9.4. The following gradient was run: 0 min: 90 % B; 14 min: 60 % B; 20 min: 95 % B; 23 min: 95 % B; 23.01 min: 90 % B, equilibration for 7 min, total run time 30 min. The triple quadrupole was operated in MRM mode, observing the following transitions: 179.0 → 89.1 and 179.0 → 59.1 for glucose; 259.0 → 96.9 and 259.0 → 78.9 for glucose-6-p; 338.9 → 97.0 and 338.9. → 79.0 for fructosebisphosphate; 169.3 → 79.0 and 169.3 → 97.0 for glyceraldehyde-3-phosphate; 89.4 → 43.1 and 89.4 → 41.0 for lactate. Analysis was performed in Skyline 64-bit (vers. 21.2.0.369).

### Isolation of pancreatic islets

For isolation of pancreatic islets, 3 ml of collagenase P (1.7 U/mg; Roche) resolved in HBSS buffer (Gibco, Thermo Fisher Scientific™) supplemented with 4.1 mmol NaHCO_3_ (Merck), was injected into the common bile duct. The pancreas was collected and incubated for 12-16 minutes (depending on the potency of the collagenase) at 37°C. The collagenase activity was stopped using HBSS (Gibco, Thermo Fisher Scientific™) containing 10% FCS and 4.1 mmol NaHCO_3_. After several washing steps using HBSS (Gibco, Thermo Fisher Scientific™) supplemented with 4.1 mmol NaHCO_3_, a density gradient centrifugation was performed using Histopaque®-1077 (Sigma-Aldrich). The enriched islets were then purified by several washing steps using HBSS. Pancreatic islets were handpicked under a microscope and used for gene expression analysis or glucose-stimulated insulin secretion.

### Glucose-stimulated insulin secretion and insulin content in isolated pancreatic islets

Pancreatic islets were cultured in RPMI media supplemented with 10 % FCS (Capricorn Scientific) and 1% penicillin/streptomycin mix (penicillin 10,000 U and 10 mg streptomycin/ml) (Sigma-Aldrich) overnight. Ten pancreatic islets were pre-incubated in Krebs-Henseleit Buffer (KHB), 0.5% BSA, and 2.8 mM glucose for 60 min at 37°C. Thereafter, islets were incubated in KHB containing 0.5% BSA, and 2.8 mM or 16.7mM glucose for 5 min (initial insulin secretion) or for 30 min at 37°C. Total insulin content was measured following acid/ethanol (0.18 mM HCl in 70 % ethanol) extraction from isolated islets. Insulin levels were determined using Ultra-Sensitive Mouse Insulin ELISA Kit (Crystal Chem).

## Data Analysis

The number of independent experiments is indicated in each figure legend along with the used statistical test and p value. For single-cell experiments, individual cells were used for analysis. Statistical analyses, including Student’s t-test and Analysis of variance (ANOVA) with Tukey post hoc test, were performed on GraphPad Prism software version 9.5.1 (GraphPad Software, San Diego, CA, USA) and Microsoft Excel (Microsoft Office 2013).

## Data availability

The data supporting the findings of this research can be obtained from the authors upon a reasonable request; please refer to the author’s contributions for details regarding specific datasets.

## Code availability

The code used in this study is available from the authors on reasonable request.

## Acknowledgments

F.M, T.E., T.M., T.P., and W.F.G. are grateful to the Austrian Science Fund (FWF) for the excellence cluster 10.55776/COE14 (MetAGE). This research was further supported by the FWF via the DKplus program 10.55776/W1226 (DK-MCD) to F.M., T.M., R.B.G., and W.F.G., 10.55776/P28529, 10.55776/I3716 (to R.M.), 10.55776/P28854, 10.55776/I3792, 10.55776/DOC130 (to T.M.), and 10.55776/I4319 (to. M.S.R.); the Austrian Research Promotion Agency (FFG) grants 864690 and 870454; the Integrative Metabolism Research Center Graz; the Austrian Infrastructure Program 2016/2017 (to T.M.), and 10.55776/COE7, 10.55776/COE17, 10.55776/FG12 and 10.55776/F73 (to R.B.G.); the Styrian Government; the City of Graz; and BioTechMed-Graz (flagship project) (to T.M.); the MEFO Graz (to W.F.G.). F.E.O., J.G. and Z.K. were doctoral fellows in the doctoral program Metabolic and Cardiovascular Disease (MCD) (FWF, 10.55776/W 1226) at the Medical University of Graz. S.v.A. and A.S. are thesis students in the excellence cluster 10.55776/COE14 (MetAGE, W.F.G.). M.H. was a fellow of the Molecular Medicine (MolMed) doctoral program at the Medical University of Graz supported by MEFO Graz. We appreciate the technical assistance from Anna Schreilechner. The SIM equipment is part of the Nikon Center of Excellence, Graz, and is supported by the Austrian infrastructure program 2013/ 2014, Nikon Austria Inc., and BioTechMed (to W.F.G.). Our thanks go to Shimadzu Austria for supporting this research through instrument access given in the Metabolomics and Bioprocess Analytics laboratory (TU Wien) (R.B.G.). For open access purposes, the authors have applied a CC BY public copyright license to any author-accepted manuscript version arising from this submission.” We thank the Medical University of Graz for financial support.

